# Predicting the distributions of *Pouteria adolfi-friederici* and *Prunus africana* iconic tree species under current and future climate change scenarios in Ethiopia

**DOI:** 10.1101/2021.08.12.456155

**Authors:** Zerihun Tadesse, Sileshi Nemomissa, Debissa Lemessa

## Abstract

The distributions of the potential adaptive ranges of iconic plant species are not yet fully known especially in regions such as Ethiopia where high climatic variability and vegetation types are found. This study was undertaken to predict the distributions of the potential suitable habitats of *Pouteria adolfi-friederici* and *Prunus africana* tree species under the current and two future climate scenarios (i.e., RCP 4.5 and RCP 8.5 in 2050 and 2070) using MaxEnt software (version: 3.4.4.). Eleven less correlated environmental variables (r<0.7) were identified and used to make the prediction models. Elevational shifts of the highly suitable habitats, effects of elevation, solar radiation and topographic position in relation to the current and future climatic scenarios on the habitats were statistically analyzed using independent t-test and linear model. Under all climate scenarios, we found a decrease in the proportion of areas of highly suitable habitats for both study species. High potentials of suitable habitats for *Pouteria adolfi-friederici* are predicted to be confined to southwest, west central and south parts of Ethiopia in fragmented moist afromontane forest patches, while it is in the southwest and west central parts of Ethiopia for *Prunus africana*. On basis of vegetation types of the country, potential suitable habitats for *Pouteria adolfi-friederici* are predicted to occur in moist evergreen forest, dry evergreen forest and grassland complex and *Combretum-Terminalia* woodland vegetation types. Whereas, moist evergreen forest, dry evergreen forest and grassland complex and riverine vegetation types are predicted to comprise potential suitable habitats for *Prunus africana,* showing considerable spatial dynamics. Overall, our results suggest that the strategies deem to design biodiversity conservation should take into account the dynamics of the suitable niches of different species under different future climate scenarios.

## Introduction

Plant species have their optimum range of climatic condition to survive in their ecological zones [1]. Among others, the environmental factors limiting the adaptive range of the plant species are altitude, temperature, moisture, and soil types [2,3]. Climate, which represents a weather condition over a wide environment for a long period, plays important role to determine the distribution of plant species in their suitable ecological regions [2]. Temperature and precipitation are the most important environmental variables determining the climatic condition of ecological regions to which plant species are naturally adapted to grow [4]. Temperature and moisture can change as a result of climate change and challenge populations of a species already well adapted to the climatic conditions of ecological regions for long period [3].

However, different studies are depicting that climate change is taking place over the world due to global warming [5–12]. Climate change is mainly caused by increasing global temperature [13–15] that is initiated mainly due to greenhouse gas (GHG) emissions from multiple sources [16–18]. Thus, a change in global temperature and precipitation [19,20] can affect the distribution of global biodiversity in the future [21–23]. Plants can change their range of distribution to respond to the climate change [24–26] and either go extinct [27] or there would be shifting out from of their current ecological niches to new ecological areas [28,29].

*Pouteria adolfi-friederici* and *Prunus africana* tree species, which belong to Sapotaceae and Rosaceae families respectively, are iconic trees occurring in remnant fragments of moist and dry afromontane forests. The species are currently existing in scattered patterns with limited number in some agricultural landscapes of Ethiopia [30–32]. *Pouteria adolfi-friederici* is a timber tree that can grow up to 50 m, having a clear straight bole usually reaching over 16 m height with narrow conical canopy in moist afromontane forests within altitudinal range of 1350–2450m a.s.l in Ethiopia [31]. Its seeds are recalcitrant and sensitive to desiccation and need to be sown while they are in fresh condition [33–35 and effective germination thereof demands optimum temperature and moisture.

The other iconic tree species that has overlapping niche with *Pouteria adolfi-friederici* is *Prunus africana. Prunus africana*, the African cherry, is a valuable timber tree growing up to 40 m high with a diameter of 1m in moist and dry afromontane forests [33]. Similar to *Pouteria adolfi-friederici*, seeds of *Prunus africana* are recalcitrant and loose viability in a short period of time when exposed to low moisture condition. The bark extract of this species is used to treat benign prostatic hyperplasia and has drawn a considerable commercial interest [36]. Therefore, these two indigenous tree species, *Pouteria adolfi-friederici* and *Prunus africana,* were selected in our study mainly for three fundamental reasons: firstly, their seeds cannot be conserved in low temperature or in cold gene bank and, thus, they can only be conserved at their natural habitats or niches; secondly, the species have remarkable ecological and economic significance; and thirdly, *Prunus africana* is assessed as vulnerable [37,38] and the populations of both species are under extensive overexploitations for various uses. Moreover, previous studies have focused on phylogeogrpahy and medicinal uses of *Prunus africana* [36,39,40] and only limited study was conducted on *Pouteria adolfi-friederici*. Afromontane forests of Ethiopia where these iconic tree species are mainly confined to have been under immense threat due to deforestation problem [41–43]. The deforestation has resulted in diminishing of suitable habitats of these important tree species although their potential distributions in Ethiopia under climate change is not yet well known. Thus, studying the potential suitable habitat distributions of the species could have informed decisions for their conservation interventions. Therefore, the aim of this study was to understand the distributions of *Pouteria adolfi-friederici* and *Prunus africana* tree species under the current climatic condition and predict their potential suitable habitats using two climate change scenarios (RCP 4.5 and RCP 8.5 in 2050 and 2070). Here, we hypothesized that, (1), there are unoccupied suitable niches under the current climatic condition for both species in ecosystems where previously no geographical detections were addressed, (2) there will be range shifts in distributions of potential suitable habitats for both species in the future beyond the moist and dry afromontane forests ecosystems, emanating from the changes in climatic conditions.

## Materials and Methods

### Study system

The study was undertaken within the geographical area of Ethiopia, which lies between 3°24’−14° 53’ N and 32° 42’−48°12’ E (Fig 1). The total area of Ethiopia is 1.13×10^6^ square km and this is approximately equal to 113,289,457 ha within the elevational range of 124 m b.s.l. to 4560 m a.s.l.

**Fig 1.**
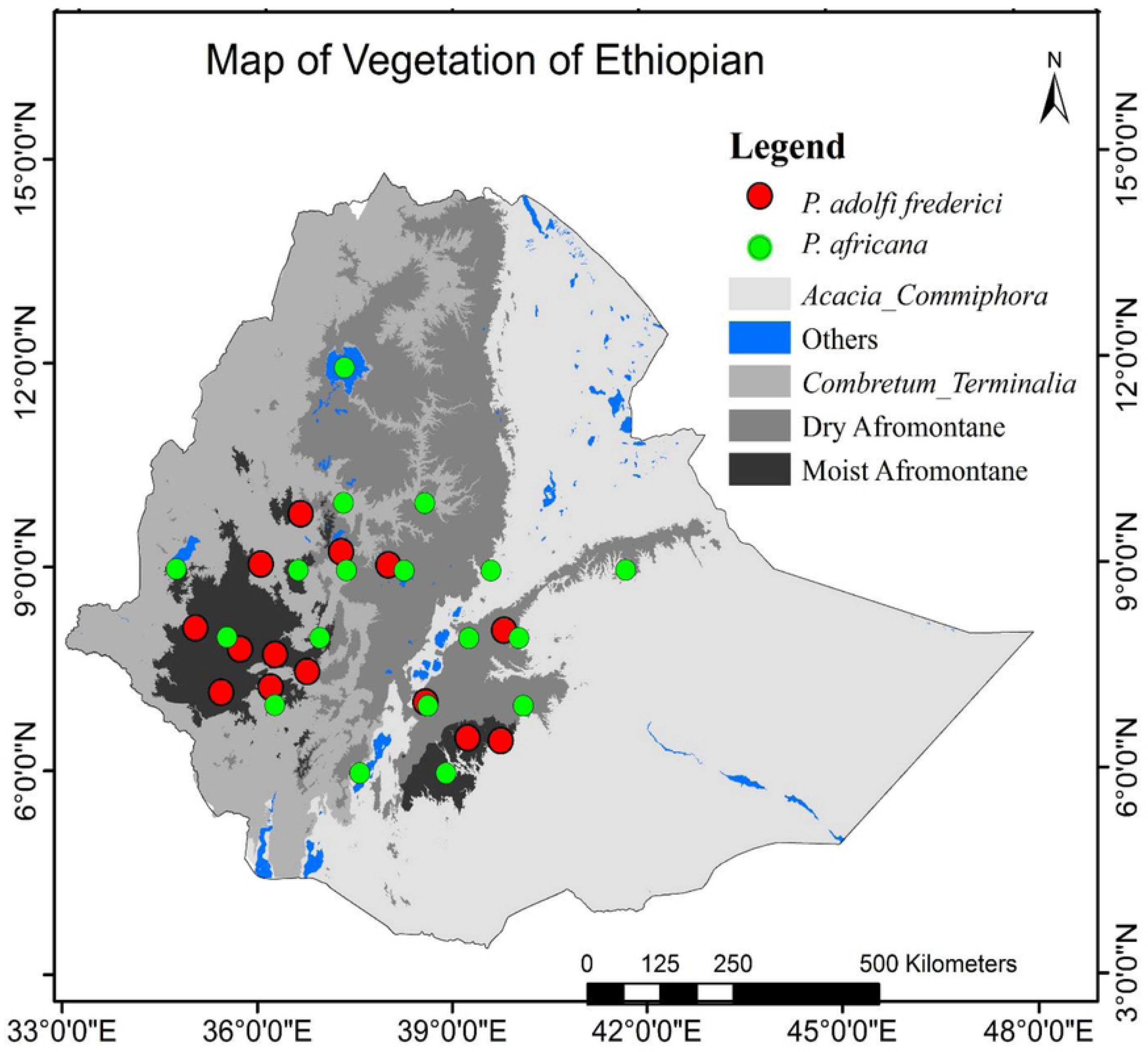
The current occurrences of *Pouteria adolfi-friederici* and *Prunus africana* tree species across five major vegetation types of Ethiopia.

Historically, the vegetation of Ethiopia has been classified in several ways, but, recently [32] has classified into 12 potential vegetation types. In the study of [44], these 12 potential vegetation types have been classified into five major vegetation groups (Fig 1 and Table 1) for conservation purposes, and we have used them for this study.

**Table 1:**
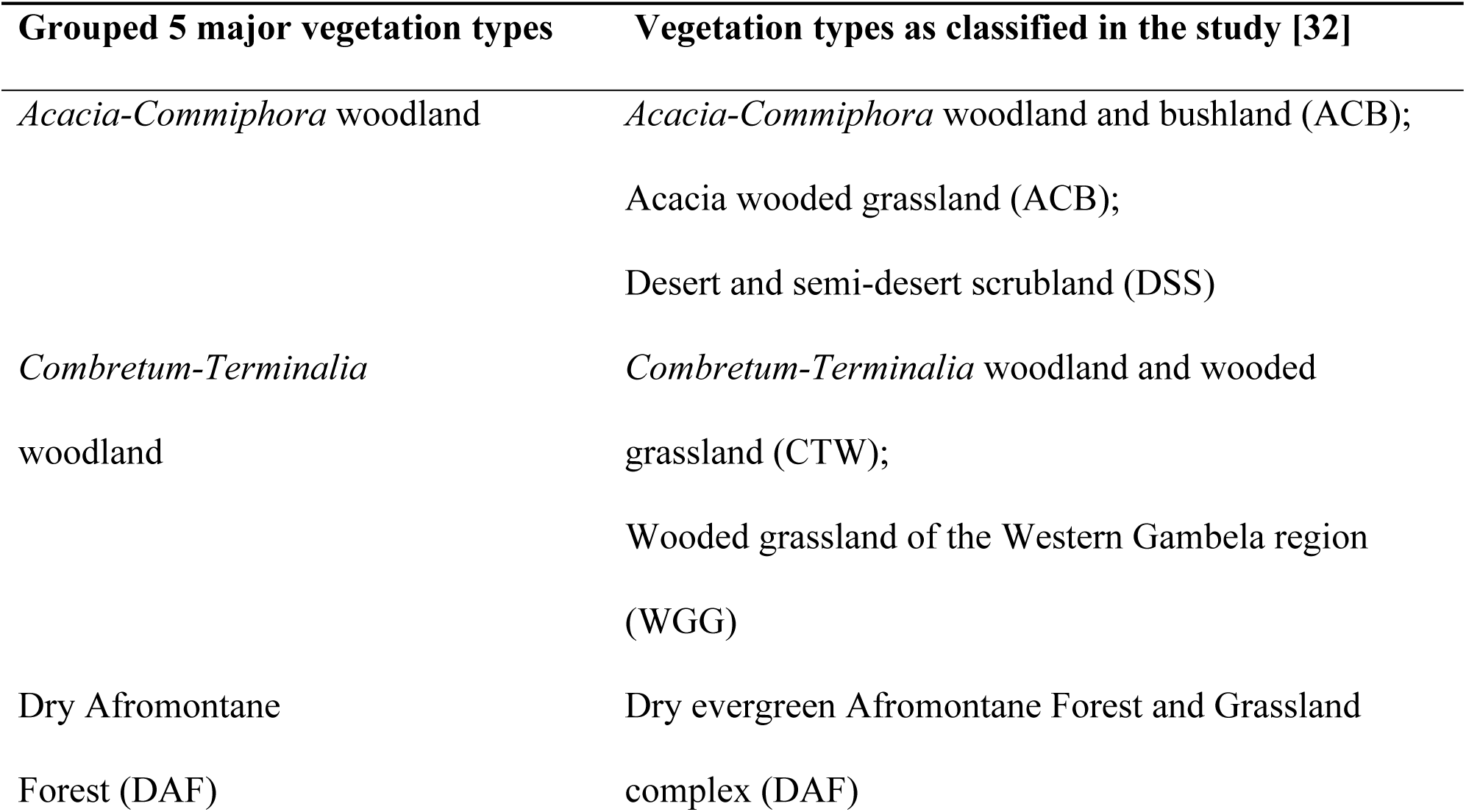
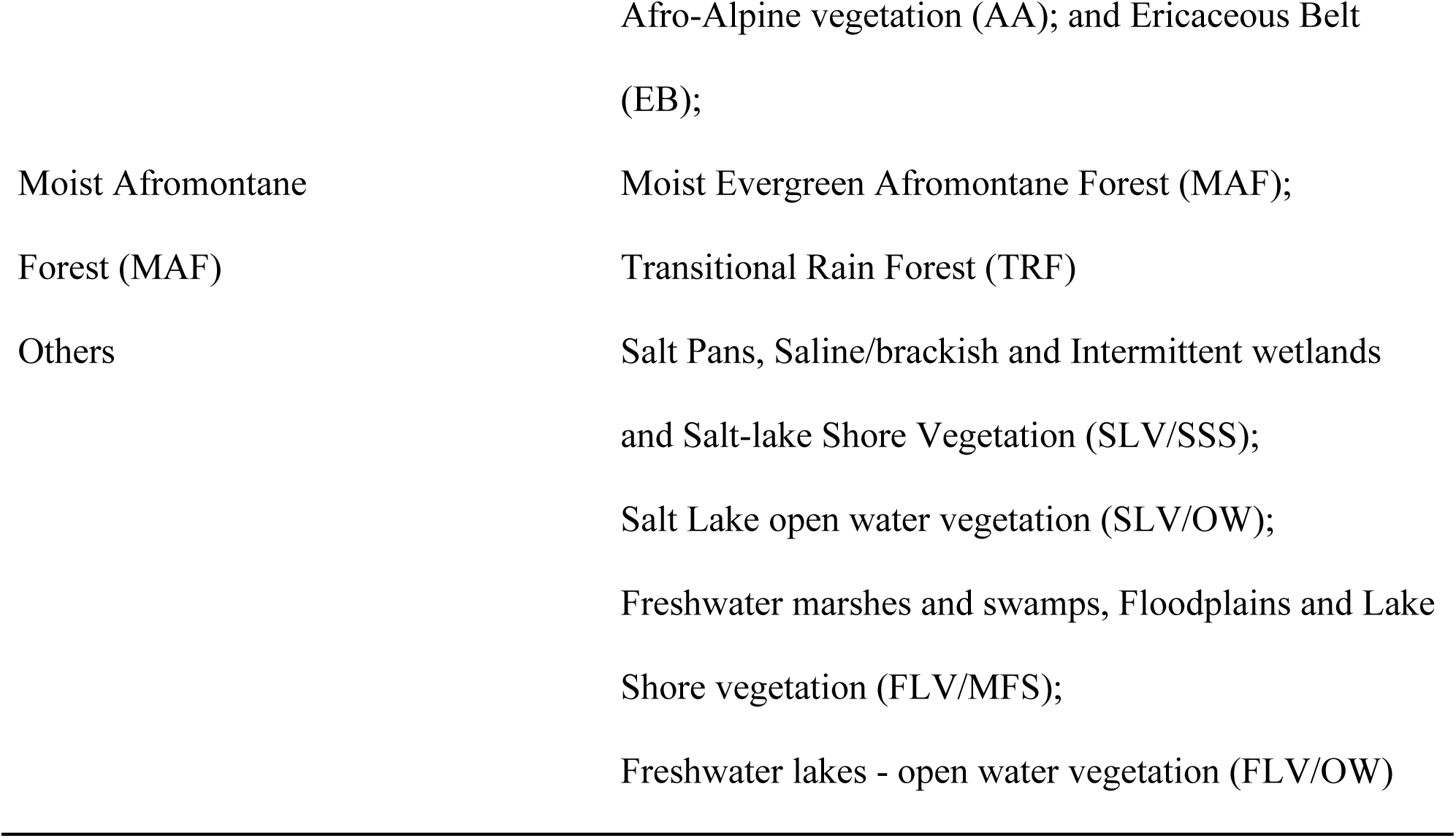
Five major vegetation groups from the 12 potential vegetation types of Ethiopia.

### Method

The species occurrences data of both species were collected from the National Herbarium of Ethiopia and from GBIF (www.gbif.org). Moreover, geographical coordinates of species presence locations were collected using a handheld Garmin GPS in west central areas of the country during a field work. In total, 94 coordinates (i.e., 30 for *Pouteria adolfi-friederici* and 64 for *Prunus africana*) points were used in this study.

As the current input data, the 19 bioclimatic variables (1970−2000) with a spatial resolution of about 1 km^2^ were downloaded from www.worldclim.org/data/worldclim21.html. Furthermore, solar radiation index (sri) was downloaded from www.worldclim.org/data/worldclim21.html while indices of topographic position and elevation data were downloaded from https://www.earthenv.org/topography.

The input data for future climate scenarios of the Coupled Model Inter-comparison Project phase (CMIP5) was used as stated in the studies [45,46]. Here, the downscaled data of Representative Pathways with intermediate (RCP 4.5) and highest concentrations (RCP 8.5) of two periods (2050 and 2070) were downloaded from http://www.worldclim.com/CMIP5_30s. These trajectories are based on their radiative forces (w/m^2^) and are expected to change at the turn of the centurey [47,48].

### Data analysis

Environmental variables were clipped into Ethiopian boundary shape file using ArcGIS (version: 10.5) and converted into ASCII format using SDMToolbox 2.4 [49]. The multicollinearity (i.e., the unnecessary correlation among the predictor variables) was checked for the current data of 19 bioclimatic variables using the pairwise correlation matrix with Pearson correlation analysis (Table in S1 Table). Eleven variables that are less correlated (r<0.7) were included in prediction models and this cut-off point was commonly applied to fit models to bioclimatic data [50–52]. The 11 bioclimatic variables used in the prediction models are mean monthly temperature, isothermality, temperature seasonality, mean temperature of coldest quarter, precipitation seasonality, precipitation of driest quarter, precipitation of warmest quarter, precipitation of coldest quarter, elevation, solar radiation index and topographic position index.

The species distribution modelling was run with MaxEnt software (version: v.3.4.4., R Core Team, 2021). The modelling was performed with a random test percentage at 25%, regularization multiplier 1, maximum background points of 10,000, maximum iterations at 5000, convergence threshold held at 0.00001 and 10 replicates with averaged results. The model was evaluated by calculating the AUC of ROC graph [53–56]. Concerned average value of AUC graph < 0.7 is bad, 0.7−0.8 is good and 0.9–1 is excellent for the species distribution modelling as explained in the study [57].

To analyze habitat suitability, the output maps of 10 replicate runs (0.0−1.0 categories) were used in ArcGIS. Accordingly, five habitat suitability classes were identified as highly suitable (0.85−1.00), suitable (0.69− <0.85), moderately suitable (0.54− <0.69), less suitable (0.38− <0.54) and unsuitable (0.0− <0.38), based on range values of band categories. The area extents were computed in hectare (ha) from the number of pixels counts for each suitable habitat classes. The change in the proportion of the habitats under current and future climate change scenarios (RCP 4.5 and RCP 8.5) was analyzed by overlying MaxEnt output maps over the shape files of the vegetation classes of Ethiopia. Predicted geographic areas in Ethiopia were extracted from Ethiogis database and the area of each was calculated using ArcGIS (version: 10.5).

### Model performance evaluation

The model outputs for both *Pouteria adolfi-friederici* and *Prunus africana* tree species indicated that all the test omission rates averaged over ten replicate runs were close to the predicted omission on the training and test rates under the current and future conditions. The average AUC for *Pouteria adolfi-friederici* showed great model fit under the current (0.926), RCP 4.5 (0.911, 0.903) and RCP 8.5 (0.901, 0.910) scenarios, at 2050 and 2070. Average AUC for *Prunus africana* also indicated very good model fit under the current (0.887), RCP 4.5 (0.868, 0.880) and RCP 8.5 (0.885, 0.896) scenarios, at the 2050 and 2070.

The differences in elevational shifts of the habitat suitability categories under current and future climatic scenarios were tested using independent t-test as implemented in R version: 4.1.0. [58]. Moreover, three environmental variables, namely, elevation, solar radiation index and topographic position index, the values of which do not vary across the climatic scenarios, were used to statistically analyze their effect on the different categories of habitat suitability using linear model in R.

## Results

### Habitat suitability mapping and modelling

The environmental variables significantly contributing to the distributions of habitat suitability of *Pouteria adolfi-friederici* were solar radiation, precipitation of warmest quarter, precipitation of driest quarter, precipitation seasonality, mean temperature of coldest quarter and elevation. Similarly, solar radiation, precipitation of warmest quarter, mean temperature of coldest quarter and mean monthly temperature significantly contribute to determine the distribution of *Prunus africana* under the current (1970−2000) and future (2050 and 2070) climate scenarios (Table 2).

**Table 2:**
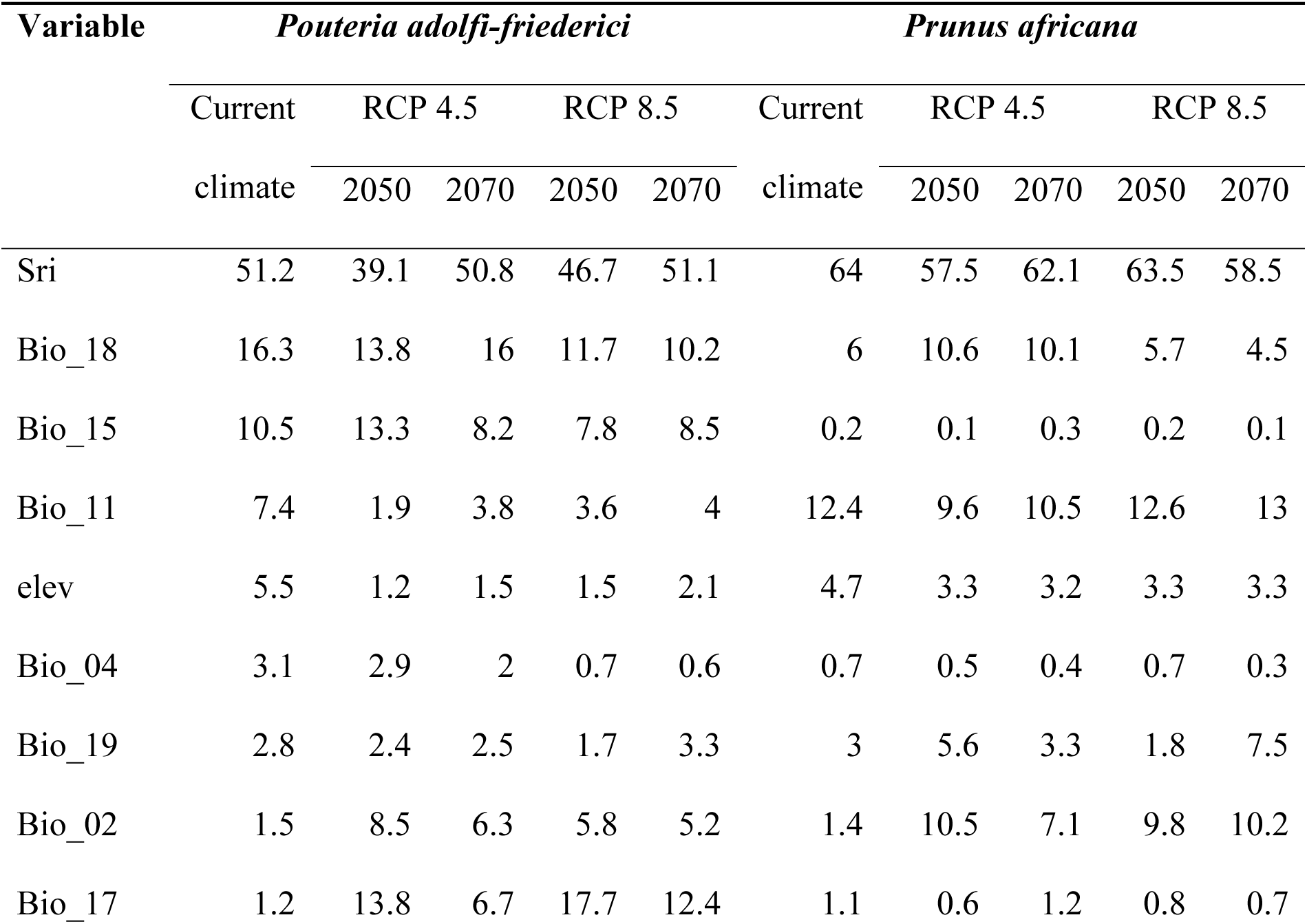
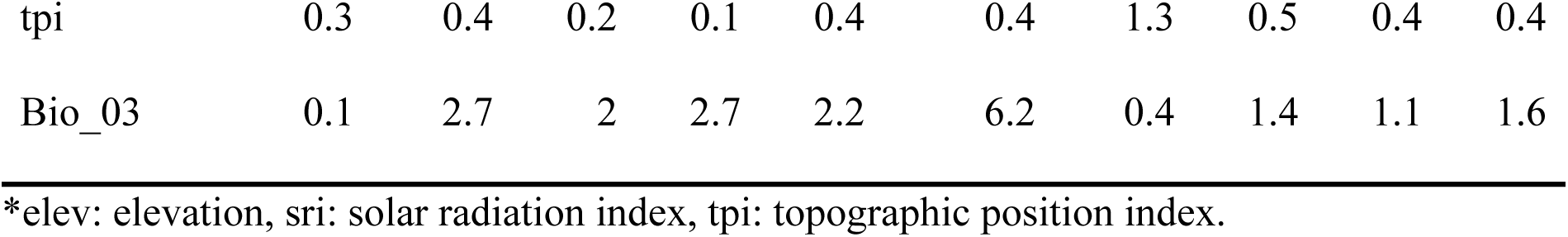
The contributions of environmental variables (%) in spatial prediction models of suitable habitats under the current and future climate scenarios of 2050 and 2070 periods.

Response curves are good informative and indicators for environmental variables contributing to the high probability of the occurrence of habitat suitability for species. The occurrence of highly suitable habitats of *Pouteria adolfi-friederici* was predicted in the range of 15,400−16,200 nm solar radiation, 30−50 mm of Precipitation Seasonality, 170−250 mm Precipitation of Driest Quarter and 500−700 mm Precipitation of Warmest Quarter (Fig 2).

**Fig 2.**
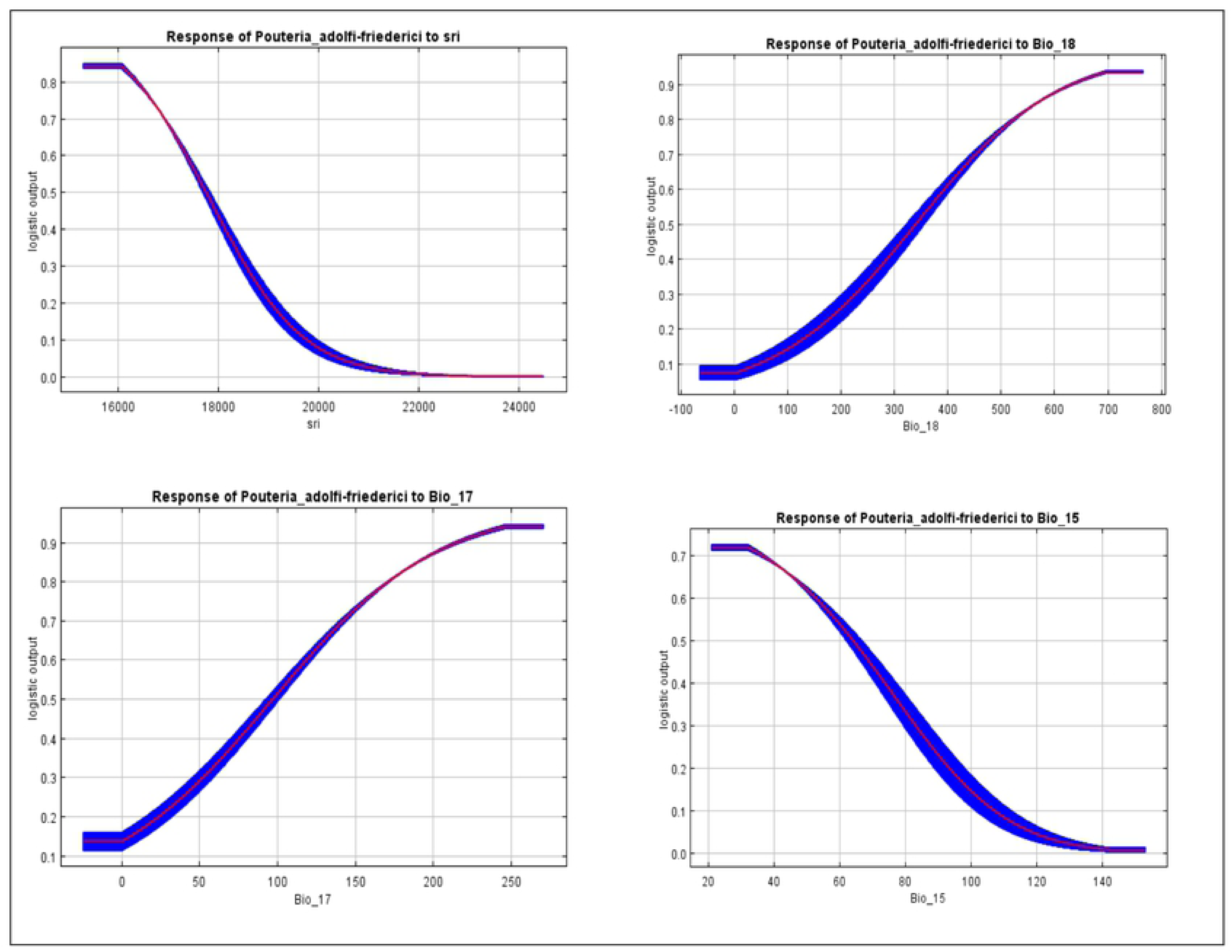
Response curves of four top contributing variables to distribution of *Prunus adofi-friederici*: solar radiation, precipitation of warmest quarter, precipitation of driest quarter and precipitation seasonality.

Similarly, highly suitable habitat for the distribution of *Prunus africana* was predicted between 15,400−16,200 nm of solar radiation, 7.5−15.5°C mean temperature of coldest quarter, 400−700 mm precipitation of warmest quarter and elevation in 2250−2700 m a.s.l. (Fig 3).

**Fig 3.**
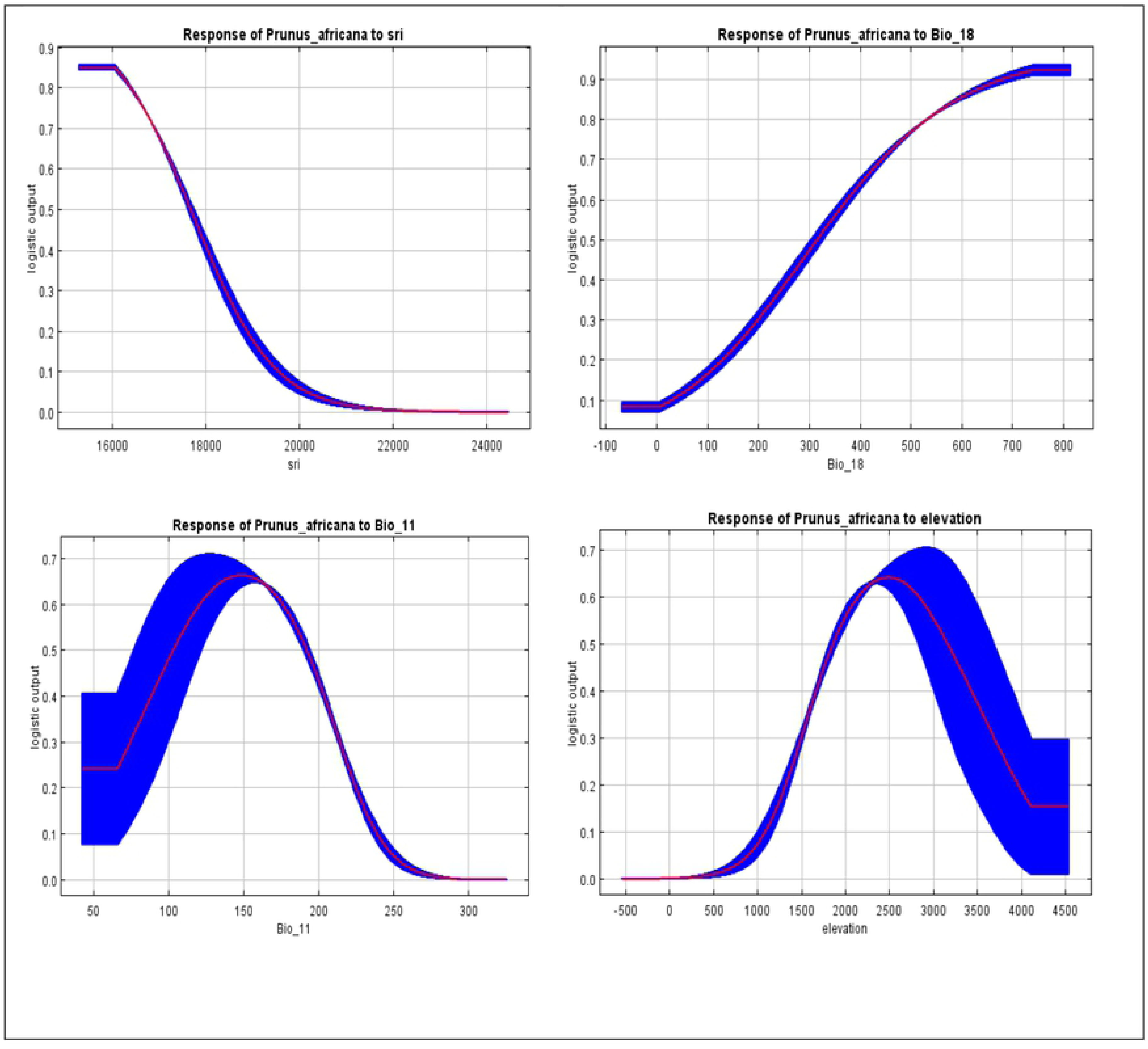
Response curves of four top contributing variables to distribution of *Prunus africana*: solar radiation, mean temperature of coldest quarter, precipitation of warmest quarter and elevation.

### Habitat suitability mapping under the current climate conditions (1970−2000)

Five habitat suitability classes were identified, i.e., highly suitable, suitable, moderately suitable, less suitable and unsuitable with different area proportions mapped for the two study species. Both moderately and less suitable habitats are areas with unfavorable conditions for both species under the current climate conditions. The distributions of highly suitable and suitable habitats of *Pouteria adolfi-friederici* are largely concentrated in southwest and with a relatively smaller amount in the southeastern part of Ethiopia (Fig 4). For *Prunus africana,* highly suitable and suitable habitats cover most of the central, eastern and western highlands of Ethiopia (Fig 5).

**Fig 4.**
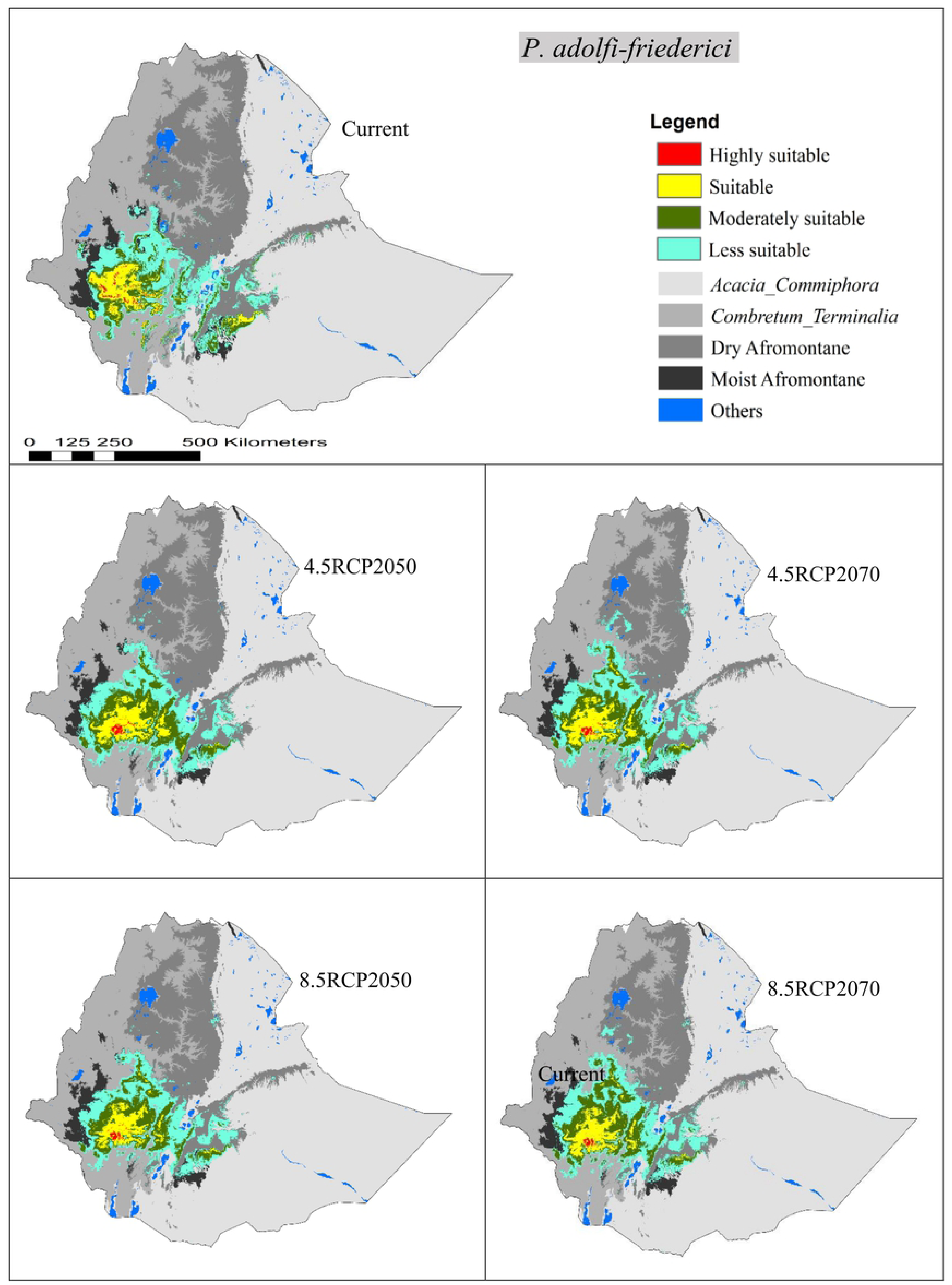
Spatial prediction models of the distributions of habitat suitability for *Pouteria adolfi-friederici* under current and future climate scenarios.

**Fig 5.**
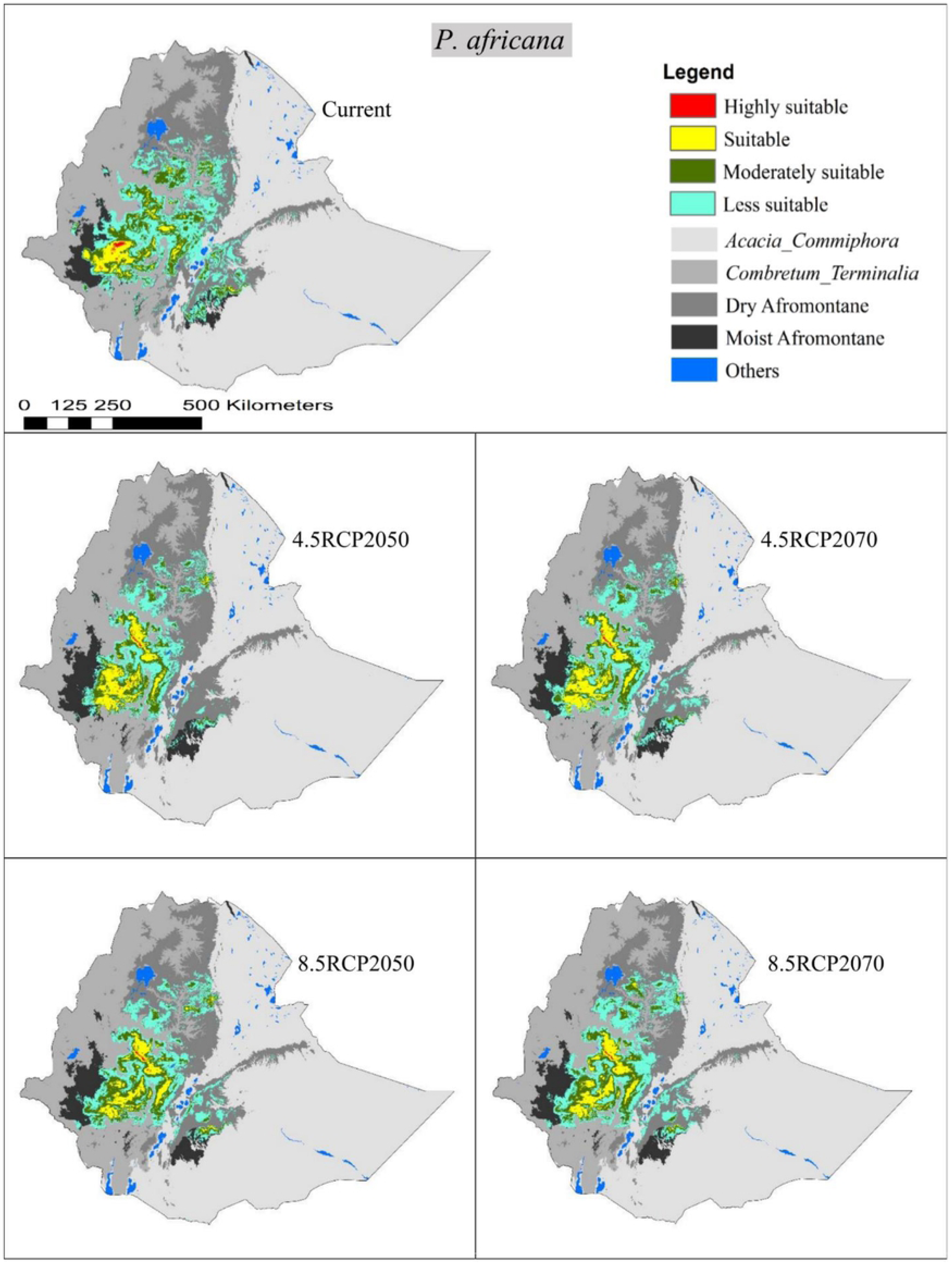
Spatial prediction models of the distributions of habitat suitability for *Prunus africana* under current and future climate scenarios.

Under the current climatic conditions, the less suitable and moderately suitable habitats cover larger areas compared to highly suitable and suitable habitats for both *Pouteria adolfi-friederici* and *Prunus africana* (Fig 6 and 7).

**Fig 6.**
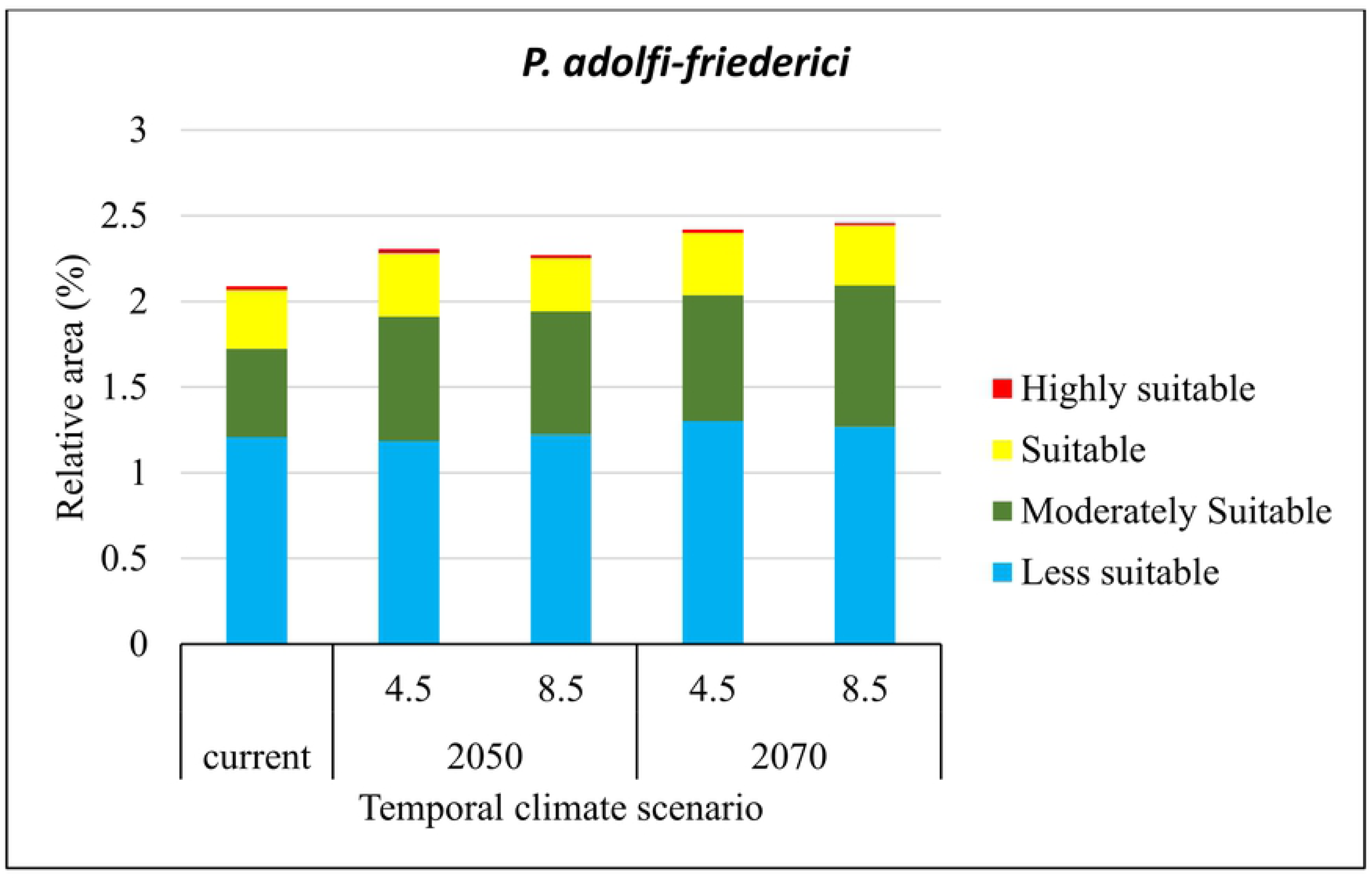
Relative area proportion (%) of suitable habitat categories of *Pouteria adolfi-friederici* under current and future climate scenarios.

**Fig 7.**
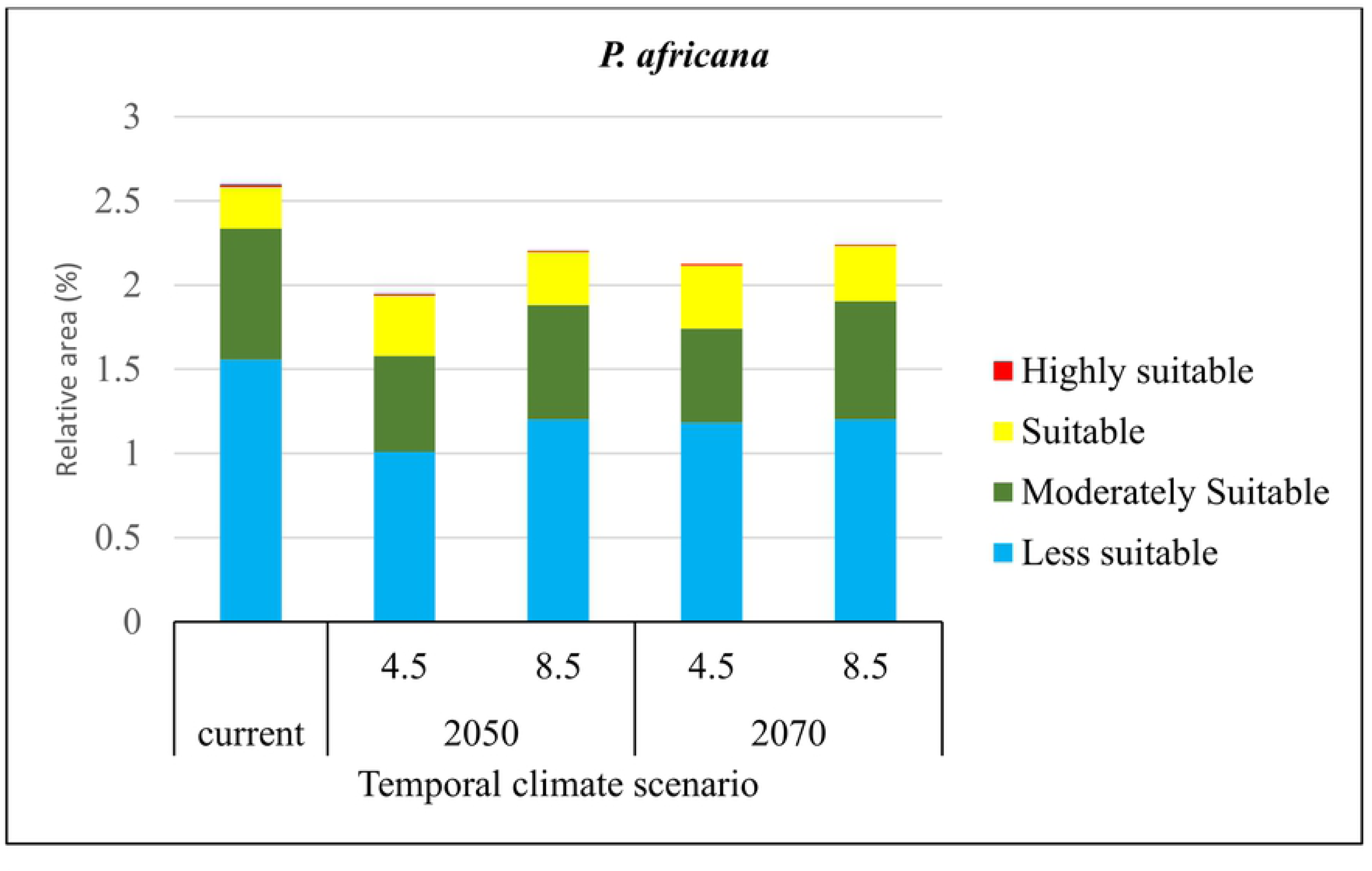
Relative area proportion (%) of suitable habitat categories of *Prunus africana* under current and future climate scenarios.

### Prediction of habitat suitability under future climate scenarios

The prediction of the distributions of habitat suitability of both study species under future climatic scenarios has shown a similar pattern to the current climate conditions. However, there are variations in terms of the percent of the relative areas across the future climate change scenarios, mainly for the three suitability classes, i.e., less suitable, moderately suitable and suitable habitats (Fig 4 and 5). Unlike these three suitability classes, the relative area of the highly suitable habitats for both tree species does not significantly vary across these scenarios (Fig 6 and 7).

### The distribution patterns of habitat suitability in the major vegetation types

#### (A) Pouteria adolfi-friederici

The relative proportion of the distribution of highly suitable habitat is higher under the current climate conditions in MAF when compared with the future climate change scenarios. However, there is a declining trend of this habitat class for this species in MAF under both future climate change scenarios (Fig 8). The proportion of suitable habitats of this species have shown a decreasing trend in all major vegetation types from the current conditions to future climate change scenarios for 2050 and 2070 but their proportions vary among the types (Fig 8). Both ACW and OTHERS (e.g. riverine vegetation) are not favorable for both species.

**Fig 8.**
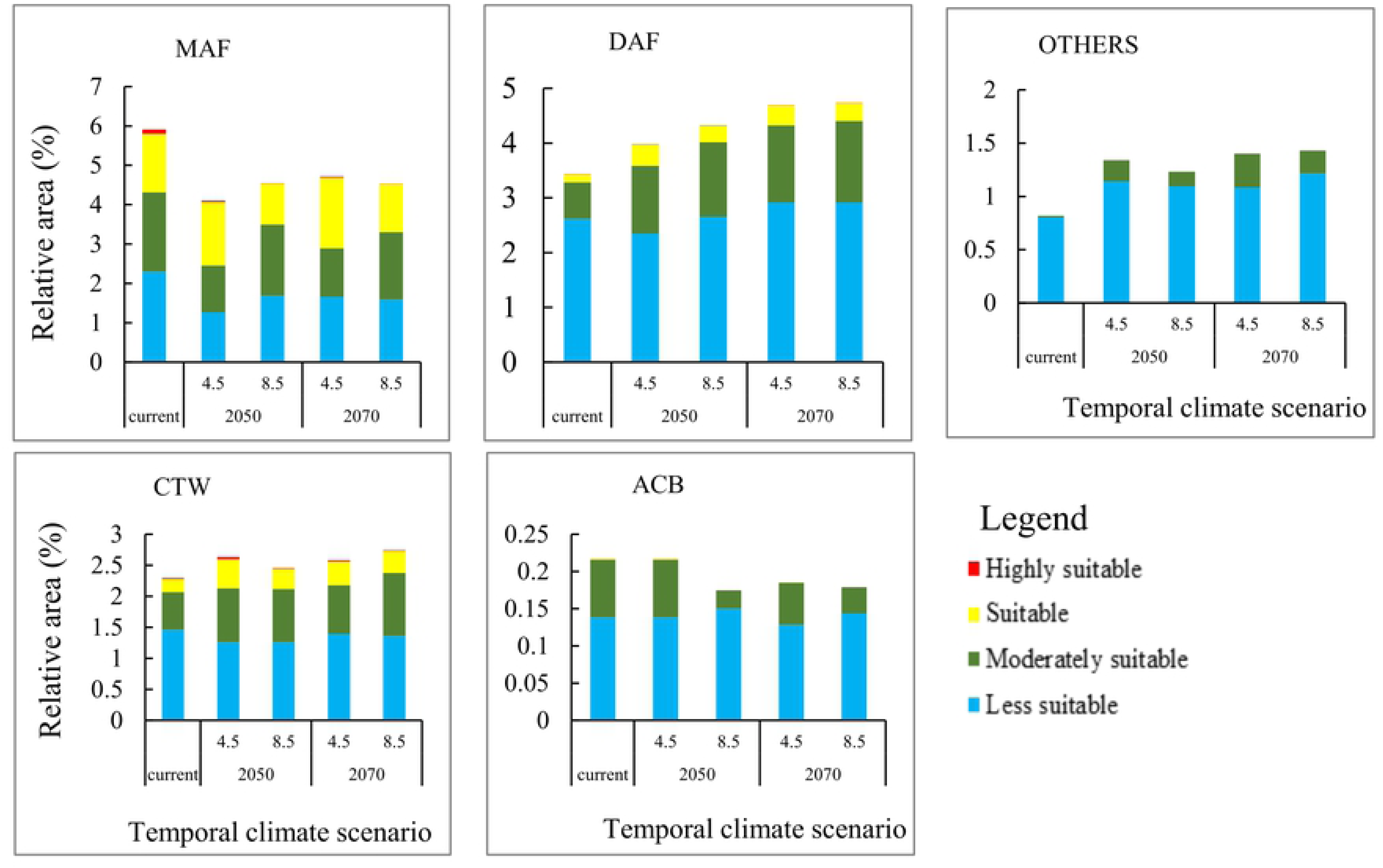
Relative area proportion (%) of suitable habitats of *Pouteria adolfi-friederici* by major vegetation types under current and future climate scenarios.

#### (B) *Prunus africana* tree species

The highly suitable habitats of *Prunus africana* tend to decline from the current to under future climate change scenarios in 2050 and 2070 (Fig 9). These habitats class will be useful for this species in CTW until 2050 for RCP 4.5, but it will be fully non-existent in 2070 and for RCP 8.5. There will also be suitable habitats in DAF for *Prunus african* despite DAF will be with less favorable habitats for this species than MAF and OTHERS. But the OTHERS vegetation type, mainly riverine, will have more favorable habitats for this species under both future climate change scenarios for 2050 and 2070 than at the current climate conditions.

**Fig 9.**
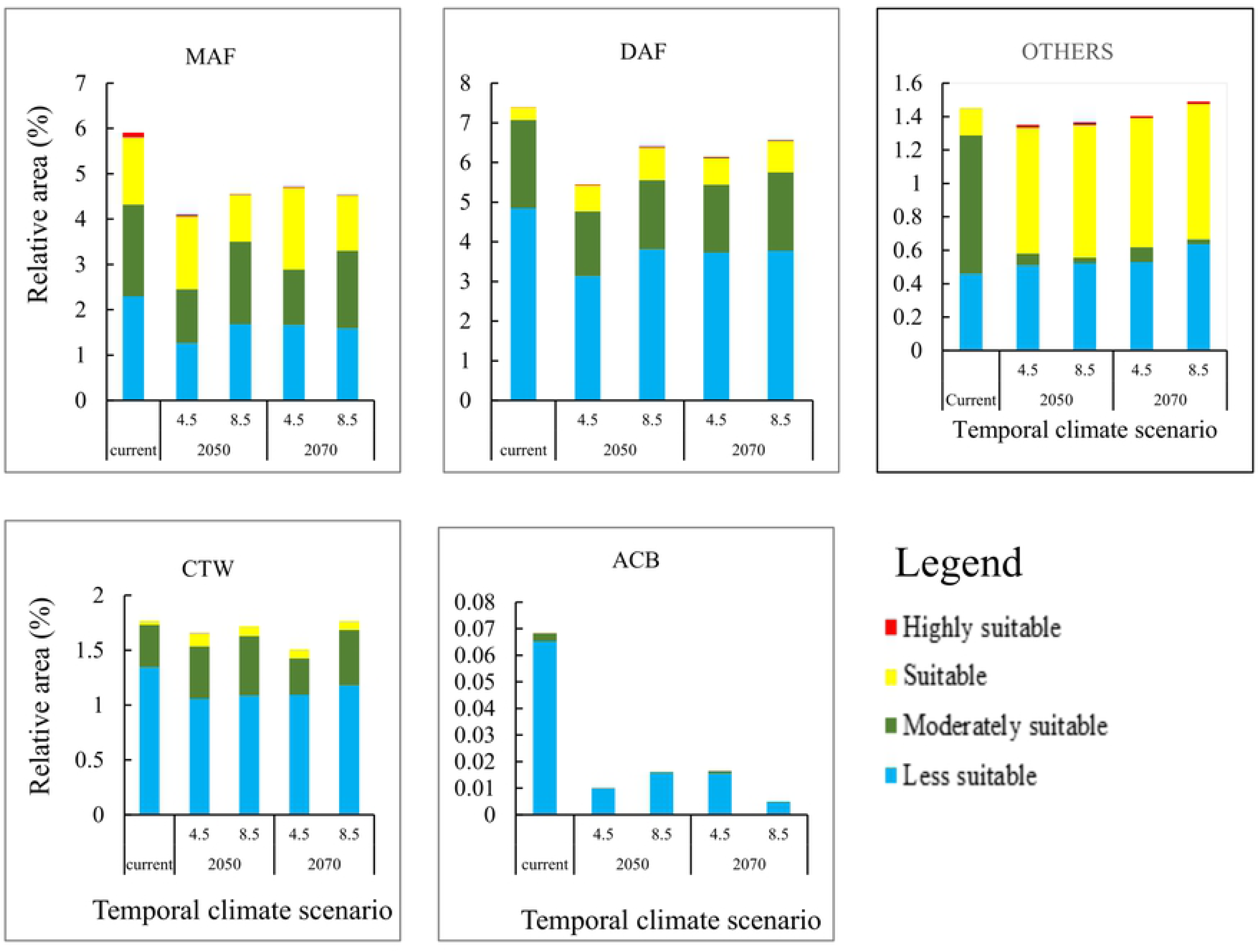
Relative area proportion (%) of suitable habitats of *Prunus africana* by major vegetation types under current and future climate scenarios.

The result of the independent t-test indicated that there is significant elevational shift in range of highly suitable habitats for *Prunus africana* between the current and future climate scenarios (Table 3) while similar trend was seen for *Pouteria adolfi-friederici* except under the RCP 4.5 at 2070, which indicated insignificant result. Moreover, the result of the linear model analyses indicated that the topographic position index has significantly affected all the three habitat categories of both study tree species (P<0.001, Table 4 and 5), and is the most influential variable to determine the distribution of potential suitable habitats for *Pouteria adolfi-friederici* and *Prunus africana*.

**Table 3:**
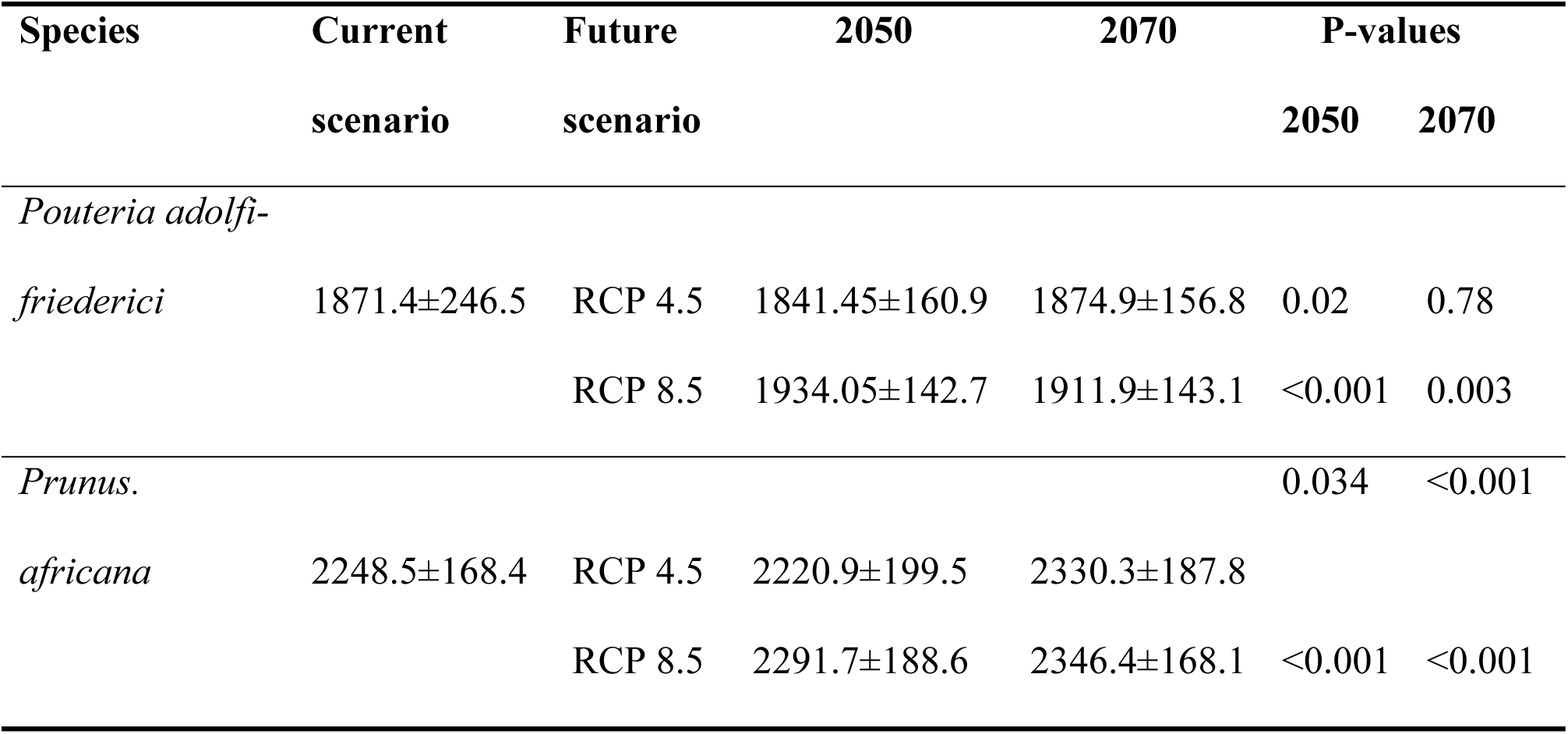
The elevational range shift status of highly suitable habitats between the current and future climate scenarios.

**Table 4.**
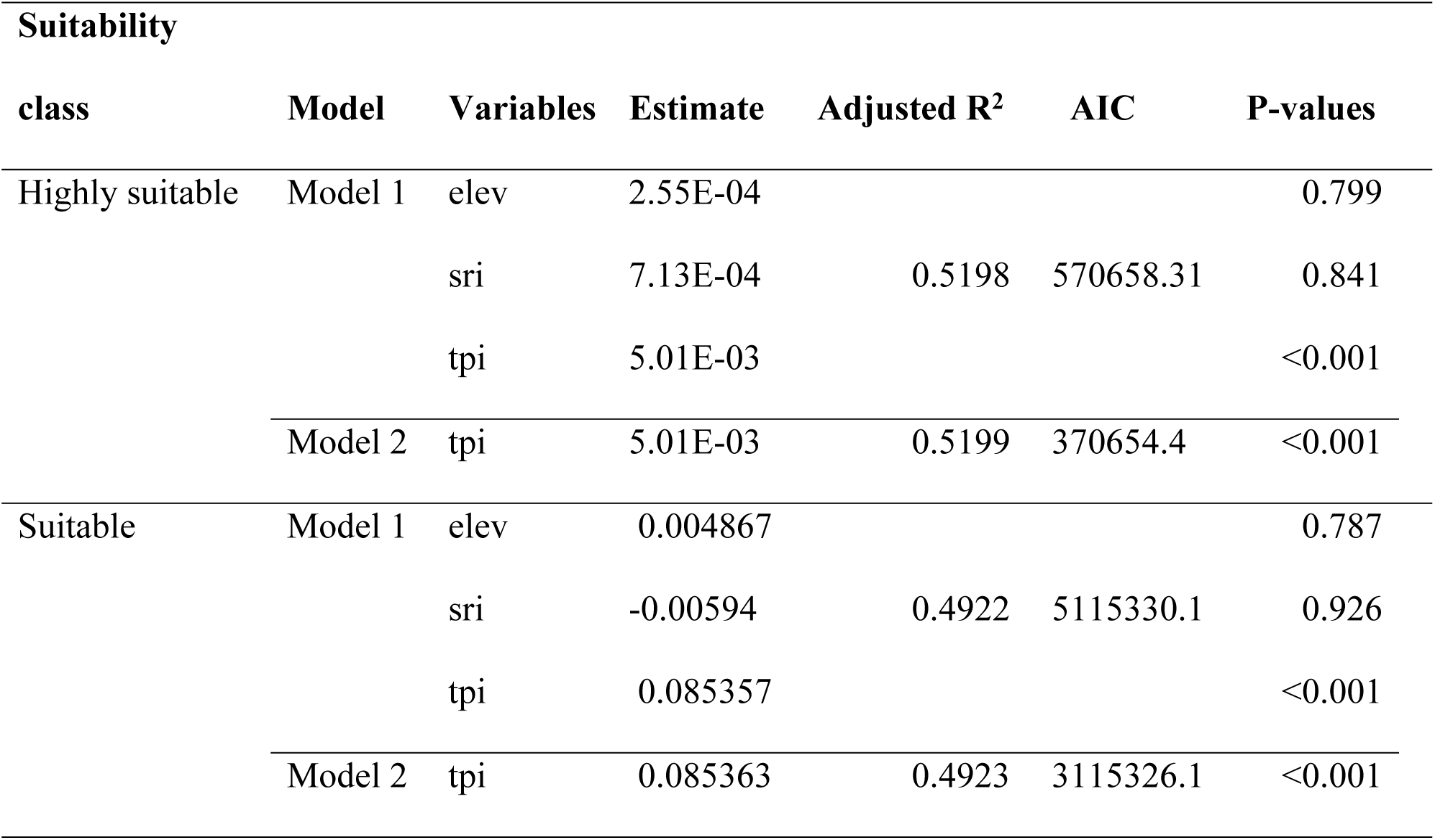
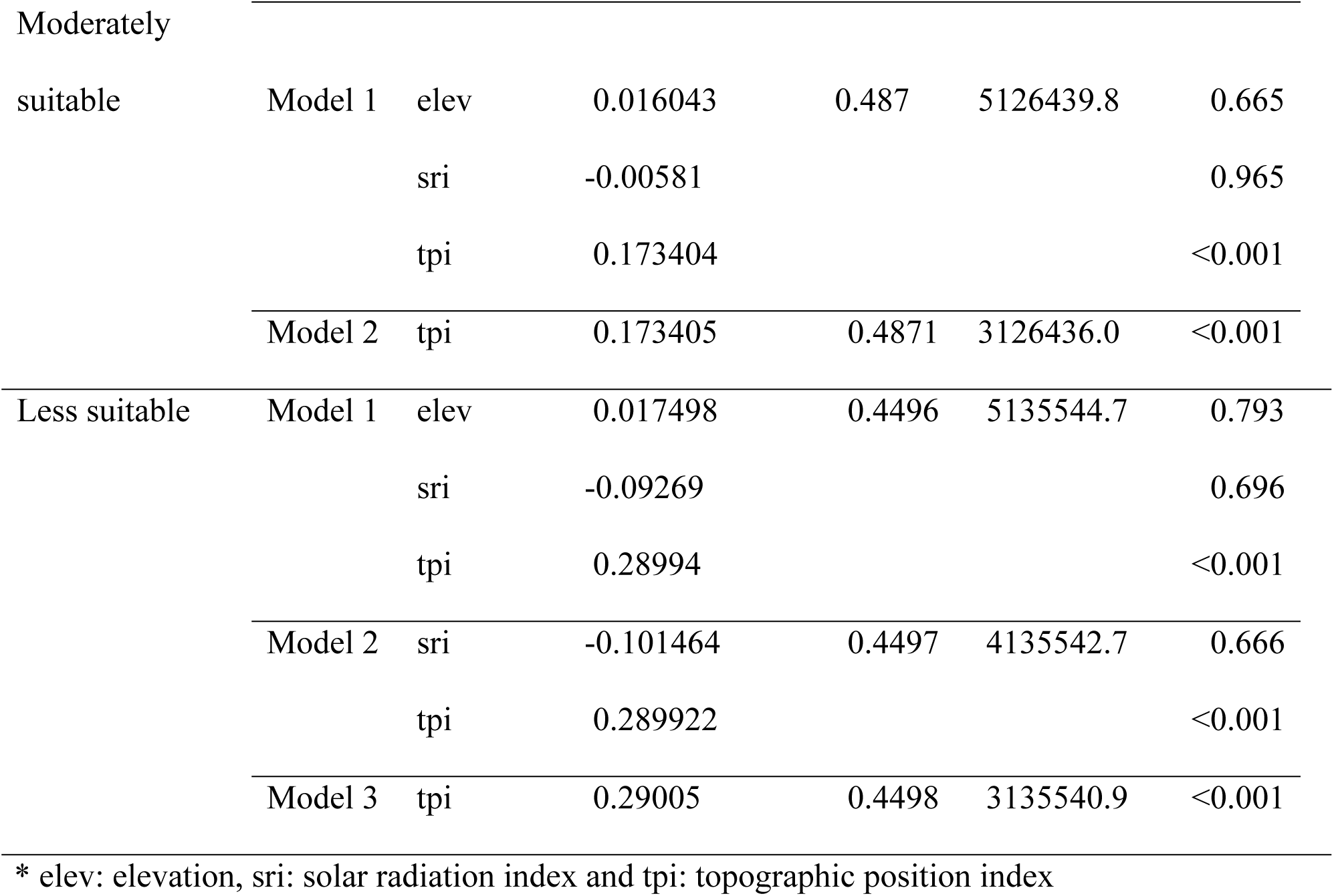
The linear models (lowest AIC and higher Adjusted R^2^) of the five habitat suitability categories for *Pouteria adolfi-friederici* under the current, 2050 and 2070 years predicted by three bioclimatic factors generated from the freely accessible Global Bioclim data.

**Table 5.**
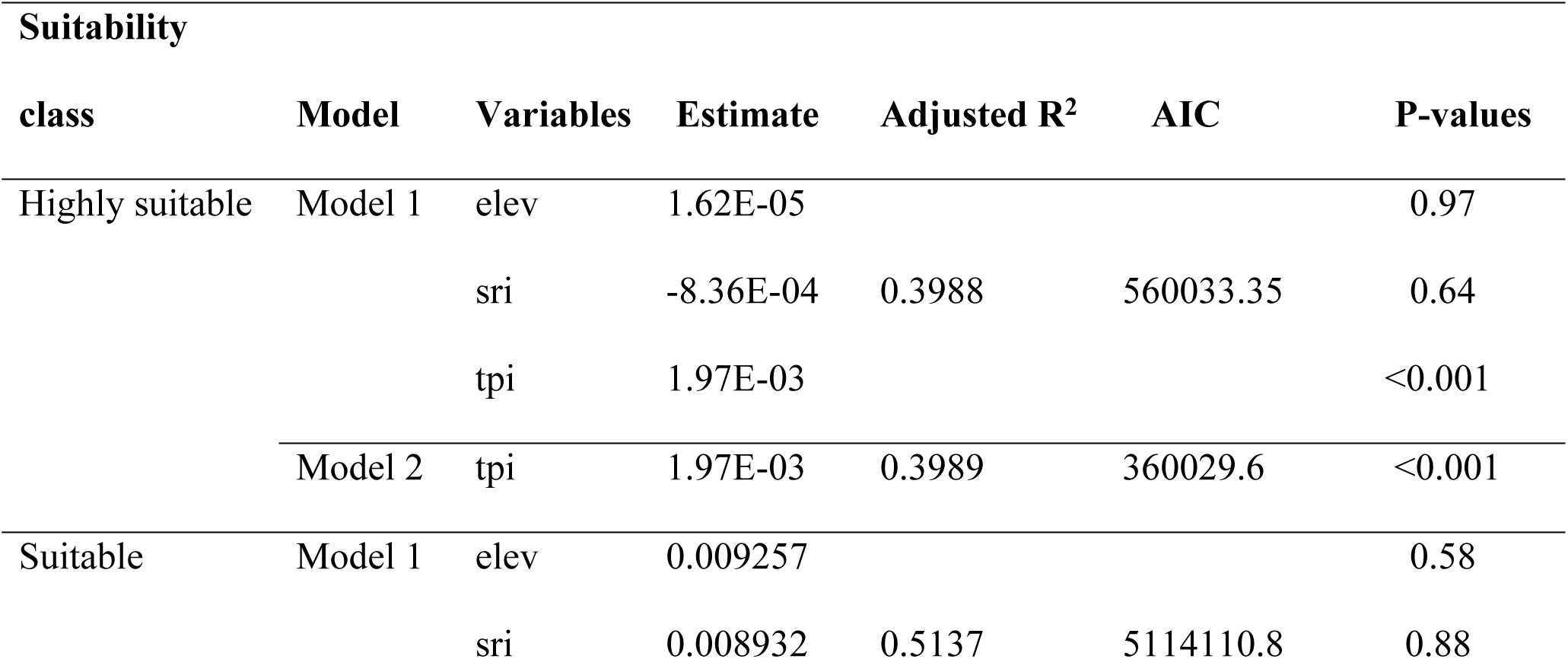
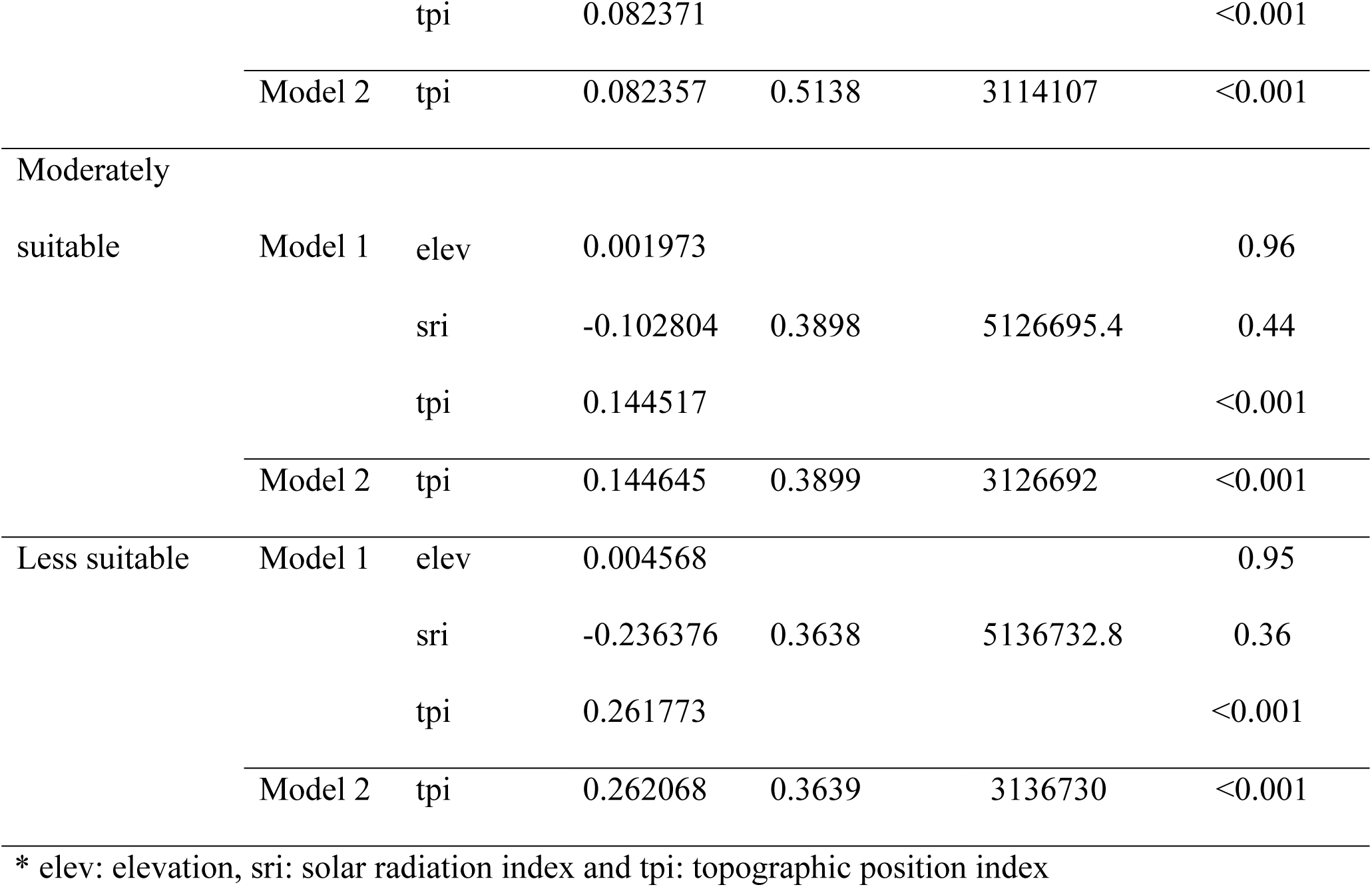
The linear models (lowest AIC and higher Adjusted R^2^) of the four habitat suitability categories for *Prunus africana* under current, 2050 and 2070 years predicted by three variables generated from the freely accessible Global Bioclim data.

## Discussions

The distributions of the species niches are reciprocally related to the dynamics in environmental variables or climatic variables. Understanding such distributions in potential suitable habitats of indigenous flora is critically important for designing conservation strategies in this era of climate change. To this end, we modeled the distributions of the potential suitable habitats of *Pouteria adolfi-friederici* and *Prunus africana* tree species under current and two future climate change scenarios (i.e., RCP 4.5 and RCP 8.5 by 2050 and 2070) and in relation to the major vegetation types of Ethiopia. Here, we show that the potential suitable habitats are predicted to be in the southwest part of Ethiopia in: moist evergreen, dry evergreen forest and grassland complex and *Combretum*-*Terminalia* woodland vegetation types for *Pouteria adolfi-friederici,* where high moisture is prevalent. Moreover, our prediction model indicated that the distribution of high suitable habitats may be in the west central parts of Ethiopia in dry evergreen forest and grassland complex and in riverine vegetation types for *Prunus africana* tree species.

Interestingly, the prediction models show that the distributions of the suitable habitats or niches of these tree species can shift among the vegetation types or ecosystems due to the effect of the future climate change scenario (Fig 4 and 5). The trend of the distributions of highly suitable habitats of both species *have* shown shifts species under different climate scenarios. In this regard, our results corroborate several previous studies that explained that climate change can affect the adaptive habitat ranges of plant species [2,50,52,59]. From the present study, such range shifts in distributions were manifested by either increasing or decreasing in areas under climate scenarios of 2050 and 2070 (Fig 6 and 7) and this trend was also highlighted with earlier findings [60,61].

Our study has revealed that the environmental factors such as, solar radiation, Precipitation of Warmest Quarter and elevation have significantly mediated the distributions of the potential suitability habitats of the two study species (Table 2) and. For example, Precipitation Seasonality and Precipitation of Driest Quarter are the drivers of the distribution of the suitable habitats of *Pouteria adolfi-friederici* while mean monthly temperature and temperature of coldest quarter are predicted to affect the habitat suitability ranges of *Prunus africana* in Ethiopia.

Understandably, such trends of fluctuations in distributions of suitable habitats may arise from the fact that the plant species have their own specific optimum demands for temperature, precipitation and altitudinal range. In this nexus, [52] from Nepal reported that the temporal variations in temperatures and precipitations are the key factors in fluctuations of the adaptive ranges of plant species. On the other hand, in the tropics, the magnitude of the solar radiation remains a constant input under both current and the future climate scenarios and as a result may not be accountable for the variations of the distributions of the potential suitable habitats of the different plant species (Table 5). Of course, the significance of the impacts of these environmental factors is related to the elevational gradients. As a result. different plant species may move out of their fundamental niches and occupy the realized ecological zones; yet, if failed to adapt may go extinct under the future climate change scenarios [62,63]. Unequivocally, several previous findings have related that change in climatic conditions alter the distributions of biological species [64–66].

Among the biological characteristics that limit the distributions of the plant species could be their dispersal mechanisms besides the impacts climatic change [2]. Thus, despite the southern parts of the southwest regions and west central areas will be highly suitable habitat potential for *Pouteria adolfi-friederici* and *Prunus africana*, the species may not fully occupy the predicted suitable habitats due to the recalcitrant nature of their seeds, which are sensitive to fluctuation of environmental variables and liable to pest and disease damages [67,68]. Moreover, their seeds may not reach the suitable areas because of limitations of appropriate dispersal agents of the species.

### Patterns of distribution of suitable habitats in relation to major vegetation types

The prediction models indicated that, under all the climate scenarios, moist afromontane, dry afromontane and *Combretum-Terminalia* woodland vegetation types were found to be highly suitable habitats of *Pouteria adolfi-friederici* and *Prunus africana* during the periods of 2050 and 2070 (Fig 8 and 9). Moreover, from other group of vegetation types, the riverine vegetation was identified to be a high suitable habitat of *Prunus africana*, which is most likely favoured by microrefugia habitat moderated by local climate condition (mainly moisture) associated with the riverine. The highly suitable areas identified for the study species within the *Combretum-Terminalia* woodland vegetation are contributed due to the presence of confined suitable habitats within the *Combretum-Terminalia* Woodland and in the ecotones that moist afromontane and dry afromontane vegetation types share with the *Combretum-Terminalia* woodland vegetation type. The highly suitable areas of *Prunus africana* in *Combretum-Terminalia* was identified for RCP 4.5 by 2050 embedded in ecotones in the transition among moist afromontane, Combretum*-Terminalia* and dry afomontane vegetation types (Fig 5). However, these habitats seem unfavorable for this species under RCP 8.5 may be due to the impact of high temperature.

The spatial distributions of potential suitable habitats of *Pouteria adolfi-friederici* and *Prunus africana* tree species showed significant shifts along elevational gradients under the current and the future scenarios (Table 3). Similarly, topographic position was found to be important factor in driving the distributions of the suitable habitats of *Pouteria adolfi-friederici* and *Prunus africana* tree species, but such impact was not found for elevation and solar radiation (Table 4 and 5).

In general, in terms of the relative proportions of the areas predicted to be suitable habitats for both tree species, the trend show that the areas are decreasing under future climate scenarios when compared with the current status (Fig 6 and 7). In this connection, our results corroborate the earlier findings that pointed out that the climate change is highly responsible for the area contraction of suitable habitats of the plant [52,69–72]. In general, it is of quite importance to undertake the prediction models of the effects of climate change on the distributions of suitable habitats of floras not only just in terms of area shift but also by ecosystems and biomes. Moreover, the present study is limited only to exploration of the impacts of the bioclimatic variables including solar radiation, elevation and topographic position indices and it may also be interesting to examine the effects of other factors such as, soil types on the distribution of floras during this changing climate under and future scenarios.

## Conclusions

The impacts of climate change and topographic factors are critical to determine the distributions of the suitable habitats for plant species. In this regard, our study denoted that the environmental factors such as, elevation, solar radiation, topographic positions, temperature and precipitation have significantly contributed in the prediction models of the distributions of the potential suitable habitats of *Pouteria adolfi-friederici* and *Prunus africana* tree species the under and future climate scenarios (RCP 4.5 and 8.5) of 2050 and 2070 periods. During these periods, the highly suitable habitats of both tree species showed spatial and elevational shifts. For example, there is a shift of highly suitable habitats of *Pouteria adolfi-friederici* from the northern and central parts of the southwest MAF to its southern areas according to the RCPs scenarios. Similarly, the highly suitable habitats of *Prunus africana* was predicted to shift from the western areas of Ethiopia to the west central regions. By sum, from our study, we conclude that the strategies wish to design national biodiversity conservation should take into account dynamics of the suitable niches of different species under changing climate scenarios for the future.

## Acknowledgements

Our thanks goes to the Plant Biology and Biodiversity Management for facilitating, and the National Herbarium of Addis Ababa University for permitting us to collect species presence data from the specimens housed in the Herbarium. Finally, we extend our heartfelt thanks to Prof. Sebsebe Demissew and Prof. Ib Friis for permitting us to use the shape file of the Potential Vegetation Types of Atlas of Ethiopia.

## Authors contributions

The authors have equal contributions to the study. They s have participated on designing the study, data collection, data analysis and interpretation, discussing the results and drawing conclusions on the findings of the study.

## Supporting Information

S1 Table. Correlation matrix among 19 current bioclimatic variables, elevation, solar radiation and topographic position indices (done using SDM_v2.4_AcrMap 10.5).

## Notes

### Competing Interest Statement

The authors have declared no competing interest.

